# Diverse crop rotations off-set yield-scaled nitrogen losses via denitrification

**DOI:** 10.1101/2025.03.15.643431

**Authors:** Aurélien Saghaï, Monique E. Smith, Giulia Vico, Samiran Banerjee, Anna Edlinger, Pablo García-Palacios, Gina Garland, Marcel G.A. van der Heijden, Chantal Herzog, Fernando T. Maestre, David S. Pescador, Laurent Philippot, Matthias C. Rillig, Sana Romdhane, Riccardo Bommarco, Sara Hallin

## Abstract

Denitrification, a major source of gaseous nitrogen (N) emissions from agricultural soils, is influenced by management. Practices promoting belowground diversity are suggested to support sustainable agriculture, but their ability to modulate gaseous N-losses via denitrification remains inconclusive. To fill this knowledge gap, we sampled 106 cereal fields spanning a 3,000 km North-South gradient across Europe and compiled 56 associated climatic, soil, microbial and management variables. We found that increased denitrification was associated with higher proportion of time with crop cover over the last ten years. Denitrification rates were best predicted by microbial biomass and microbial functional guilds involved in N cycling, in particular denitrification. We also show that several diversification practices affect the variation in denitrification predictors, suggesting a trade-off between agricultural diversification and gaseous N-losses via denitrification. However, increased crop diversity in rotations improved yield-scaled denitrification, highlighting the potential of this practice to minimize N losses while contributing to sustainable food production.

---

Globally, there is a surplus of reactive nitrogen (N) in the environment due to the widespread use of fertilizers in agriculture^1^. Nitrogen-use efficiency in cropping systems is low, with only half of the N inputs recovered in the harvested crop^2^, and the loss of N from fertilized soils represents a major threat to the integrity of both terrestrial and aquatic ecosystems^3^. Up to 200 kg N ha^−1^ year^−1^ can be lost in the form of dinitrogen gas (N_2_) and the stratospheric ozone-depleting and greenhouse gas nitrous oxide (N_2_O)^4^, but depending on the conditions, losses can be substantially higher^5^. Denitrification, an anaerobic microbial respiratory pathway, is the main source of these gaseous losses^6^, with N_2_O emissions from fertilized soils accounting for nearly half of the anthropogenic N_2_O production worldwide^7^. Thus, the reactive N surplus is also driving climate change, which could create undefined feedbacks on terrestrial N cycling^8,9^. Identifying and promoting management practices that minimize gaseous N losses is therefore critical for sustainable agricultural intensification^10^ and climate change mitigation^11^.

Cropping systems with more diverse crop rotations and longer periods with crop cover, as well as management practices such as reduced tillage and the application of organic amendments, have been advocated to increase belowground biodiversity and promote soil multifunctionality^12–14^. Diverse cropping can also provide higher grain yields^15^ and support a variety of ecosystem services^16^. Although the abundance, composition and activity of denitrifying microbial communities are influenced by agricultural management practices^17,18^, most studies cover a limited range of environmental conditions with only a few variables measured and the relative importance of management versus other environmental drivers on denitrification remains unclear^19^. Characterising the underlying factors explaining the effects of agricultural management practices on denitrification is thus key to develop effective strategies aimed at minimizing gaseous N losses through denitrification in cropping systems.

Building on our previous study conducted along a 3,000 km North-South gradient across Europe, where we showed that crop cover promotes soil multifunctionality^12^, we investigate here whether a trade-off exists between N losses via denitrification and various diversification practices, including long-term phylogenetic and functional crop diversification, the proportion of time with crop cover, tillage frequency and the application of organic and mineral fertilizers. For this purpose, we used a dataset comprising 56 climatic, soil, microbial and management variables associated to 106 soils along the European gradient (**Fig. 1**). We hypothesized that management practices enhancing belowground diversity would lead to a greater abundance of denitrifiers, as these facultative anaerobes make up a significant portion of the overall microbial community^20^. This, in turn, would increase the potential for denitrification activity, measured here as the potential production of N_2_ and N_2_O under standardized conditions using a common-garden approach (hereafter, denitrification). We then expressed denitrification in relation to the harvested yield (yield-scaled denitrification, hereafter, _y_denitrification) to determine whether any management practice can off-set denitrification activity with higher yields. Finally, we identified the best predictors of denitrification and _y_denitrification from our dataset and assessed how much of the variation in denitrification predictors could be explained by management.

**Fig. 1.**
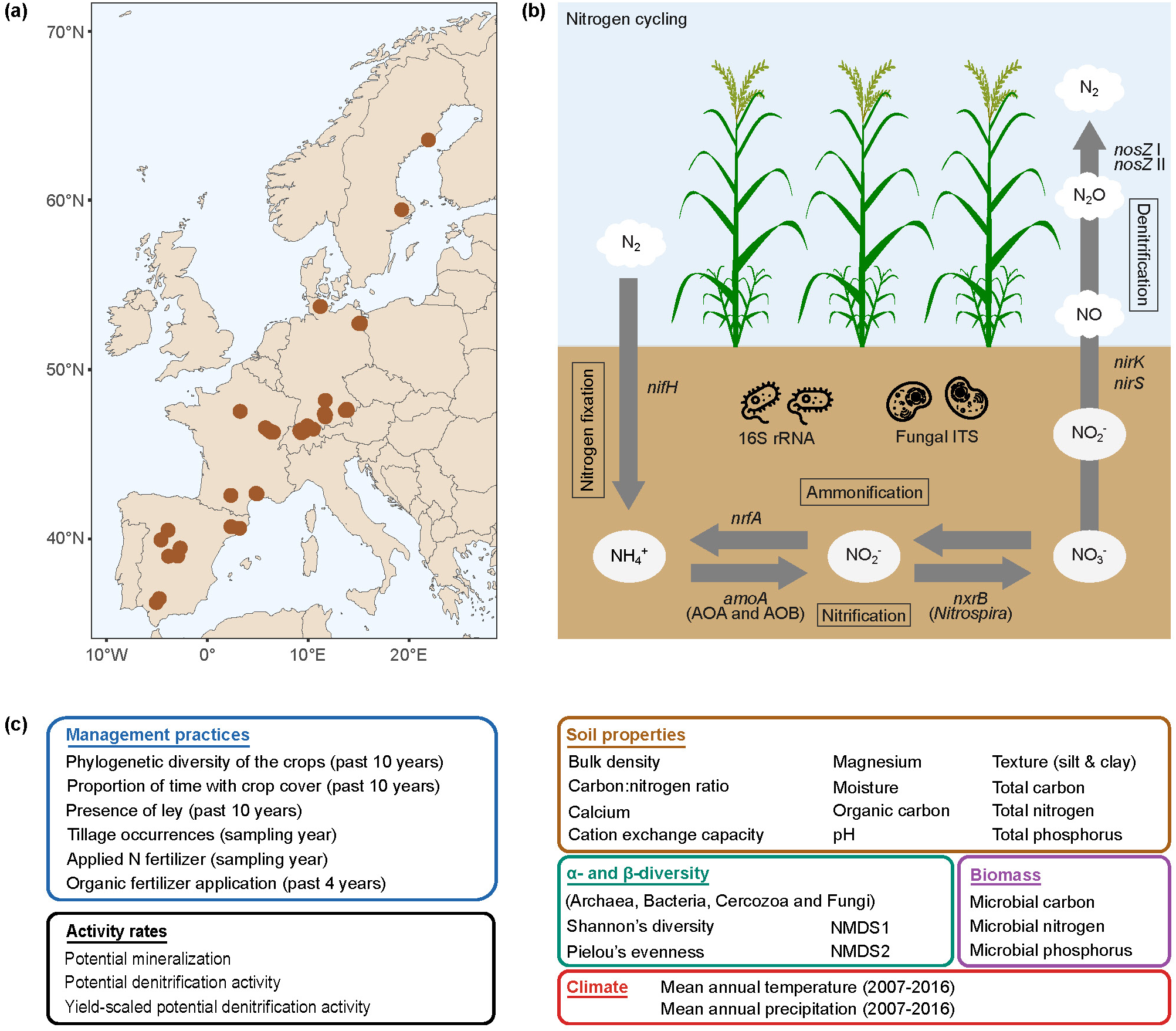
Schematics of the study design and variables included in the analyses. (a) Map of Europe showing sampling sites. (b) The nitrogen cycle with genes encoding the respective enzymes that catalyze the reactions and that were used in this study. In the nitrification process, *amoA* genes in both ammonia-oxidizing archaea (AOA) and bacteria (AOB) were quantified. For the abundance of prokaryotic and fungal communities, we quantified the 16S rRNA gene and the ITS2 region, respectively. Products and a selection of intermediates are shown by their chemical names. (c) Additional variables included were associated with management, climate, soil properties and microbial diversity, biomass, and activity. See **Table S1** for units.

## RESULTS

### Effects of management practices on denitrification and _y_denitrification

The generalized additive models including the six management practices as explanatory variables (**Figs. 2** and **3**) explained 47.3 % and 37.6 % of the deviance for denitrification and _y_denitrification, respectively. We found a non-linear relationship between the proportion of time with crop cover during the last 10 years and denitrification (**Fig. 2b**). Denitrification was highest in soils at 100 % time under cover, meaning crops were grown all year-round (including cover crops), and lowest at 50-60% time under crop cover. By contrast, lower _y_denitrification, or higher efficiency, was associated with higher phylogenetic diversity of the crops in the rotations (**Fig. 3a**). Reduced tillage was also associated with lower _y_denitrification (**Fig. 3f**), but none of the other management practices, including N-fertilization rates in the sampling year, affected denitrification or _y_denitrification (**Figs. 2** and **3**).

**Fig. 2.**
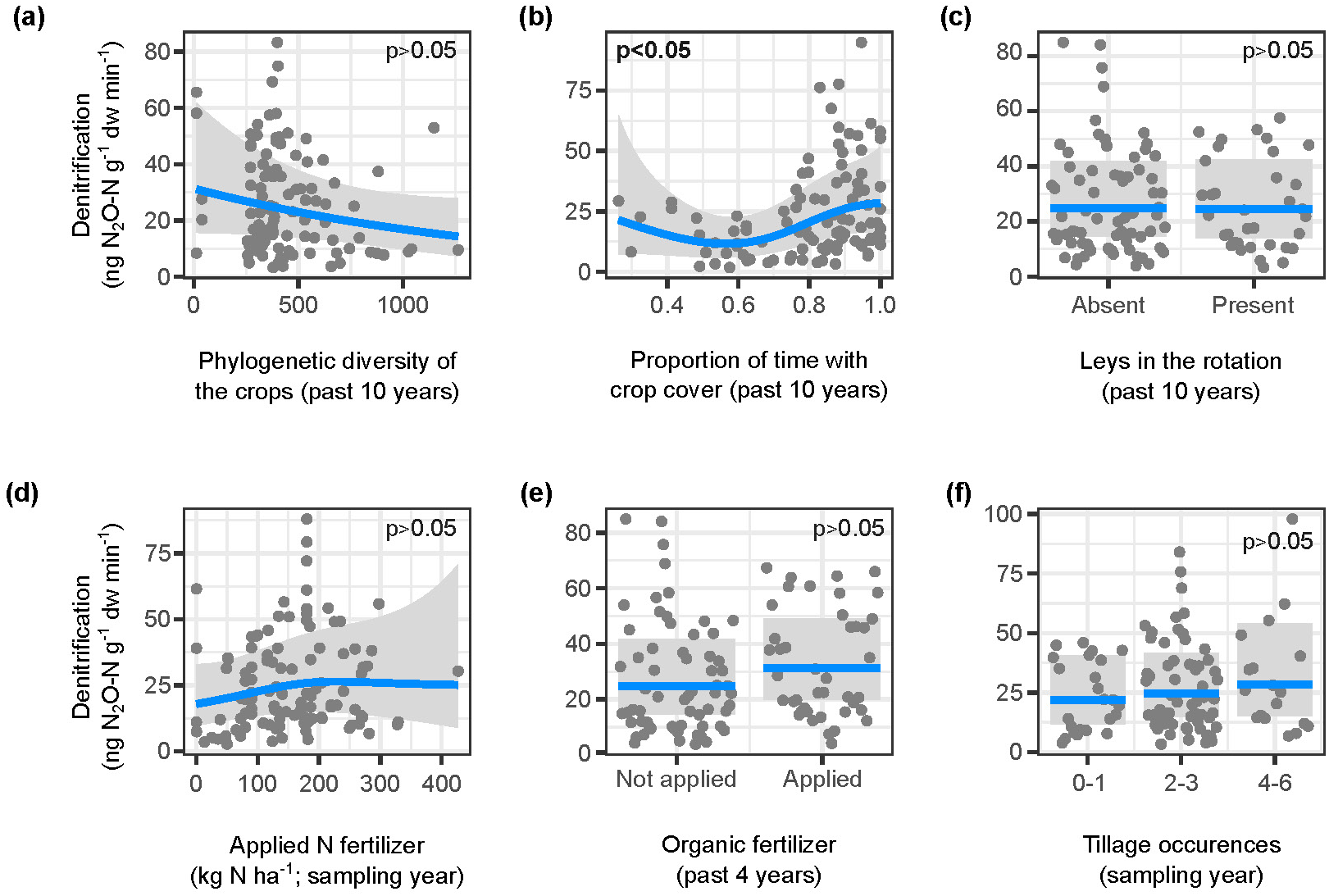
Responses of potential denitrification activity to different management practices. Responses to (a-c) phylogenetic diversity of the crops, proportion of time with crop cover and presence of ley in the rotation during the last 10 years, (d-e) amount of applied nitrogen (N) fertilizer the year of sampling (including both organic and inorganic N) and application of organic fertilizers during the last 4 years and (f) number of tillage occurrences the year of sampling. The blue line represents the estimated effect of each variable conditional upon the other terms in the model. The grey dots denote the partial residuals, and the shaded area indicates 95% confidence intervals. See **Table S2** for summary of the model statistics. P-values are indicated on the corresponding plots.

**Fig. 3.**
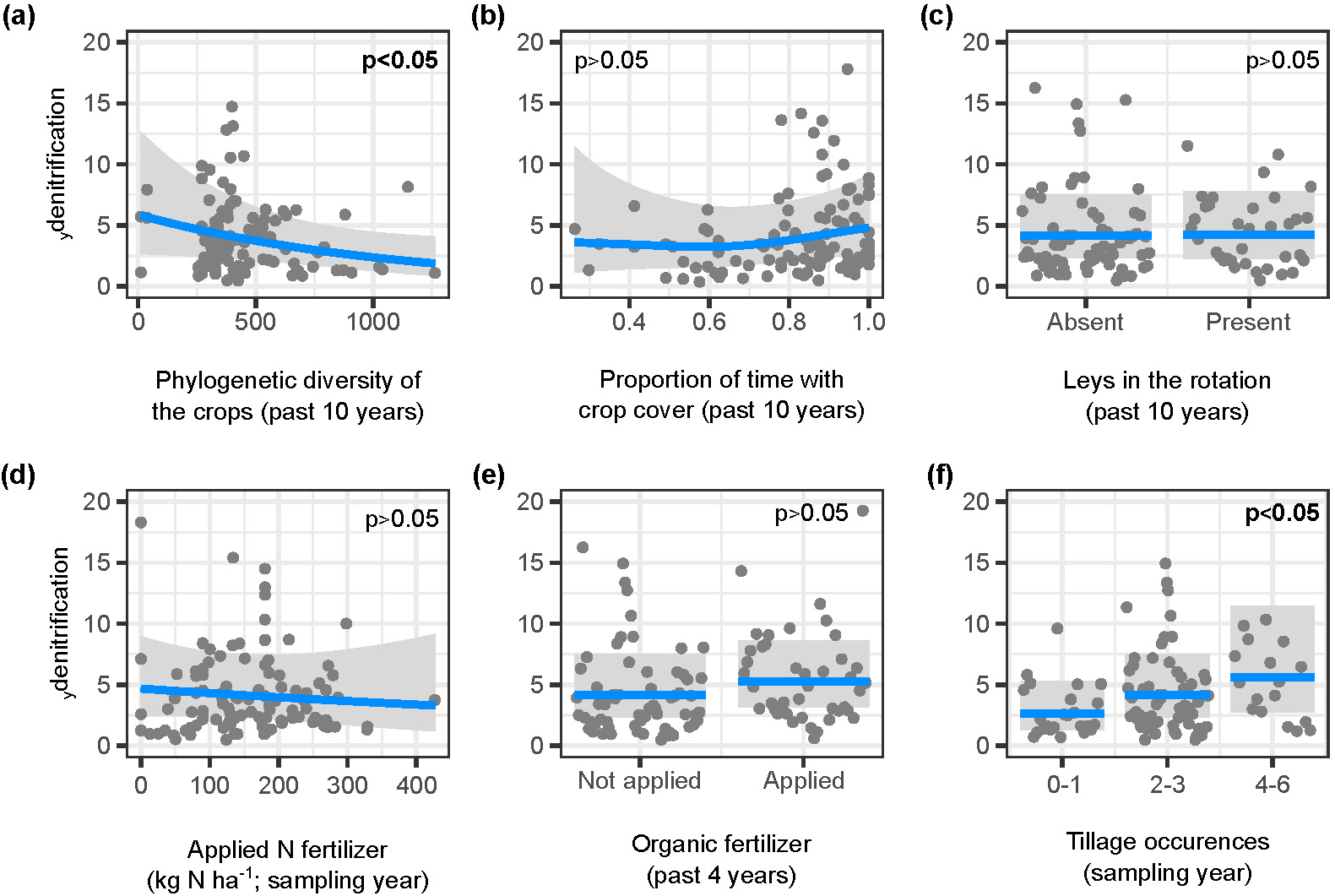
Responses of yield-scaled potential denitrification activity (_y_denitrification = denitrification/raw yield) to different management practices. Responses to (a-c) phylogenetic diversity of the crops, proportion of time with crop cover and presence of ley in the rotation during the last 10 years, (d-e) amount of applied nitrogen (N) fertilizer the year of sampling (including both organic and inorganic N) and application of organic fertilizers during the last 4 years and (f) number of tillage occurrences the year of sampling. The blue line represents the estimated effect of each variable conditional upon the other terms in the model, the grey dots denote the partial residuals, and the shaded area indicates 95% confidence intervals. See **Table S3** for summary of the model statistics. P-values are indicated on the corresponding plots.

### Predictors of denitrification activity

We then examined the relative importance of climatic, microbial, soil and management variables to predict denitrification and _y_denitrification in our dataset (**Figs. 1b-c**). Ten predictors of denitrification were retained in the final random forest model, all displaying non-linear relationships with denitrification (**Fig. 4**). The majority of predictors were microbial, with several related to the overall size of the microbial communities, as reflected by the strong positive effect of microbial biomass, the abundance of several functional guilds involved in N cycling, more specifically denitrifiers (*nirS,* nosZ clade I) and bacteria performing nitrate ammonification (*nrfA*), as well as the composition of the microbial communities (**Fig. 4b-d**). Soil predictors, and in particular total N in the range 0-3000 mg N kg^−1^ soil, were also positively associated with denitrification activity, but to a lower extent than microbial predictors (**Fig. 4a**). Regarding _y_denitrification, the final random forest model comprised a subset of the best predictors of denitrification. Higher _y_denitrification, or higher gaseous N losses in relation to the harvested yield, was mainly associated with increasing total N, particularly in the range 1500-3000 mg N kg^−1^ soil, microbial biomass and abundance of *nirS*-denitrifiers (**Fig. 5**). Neither climatic variables (mean annual temperature and mean annual precipitation over the past 10 years) nor management practices were selected as important predictors of denitrification or _y_denitrification by the random forest models.

**Fig. 4.**
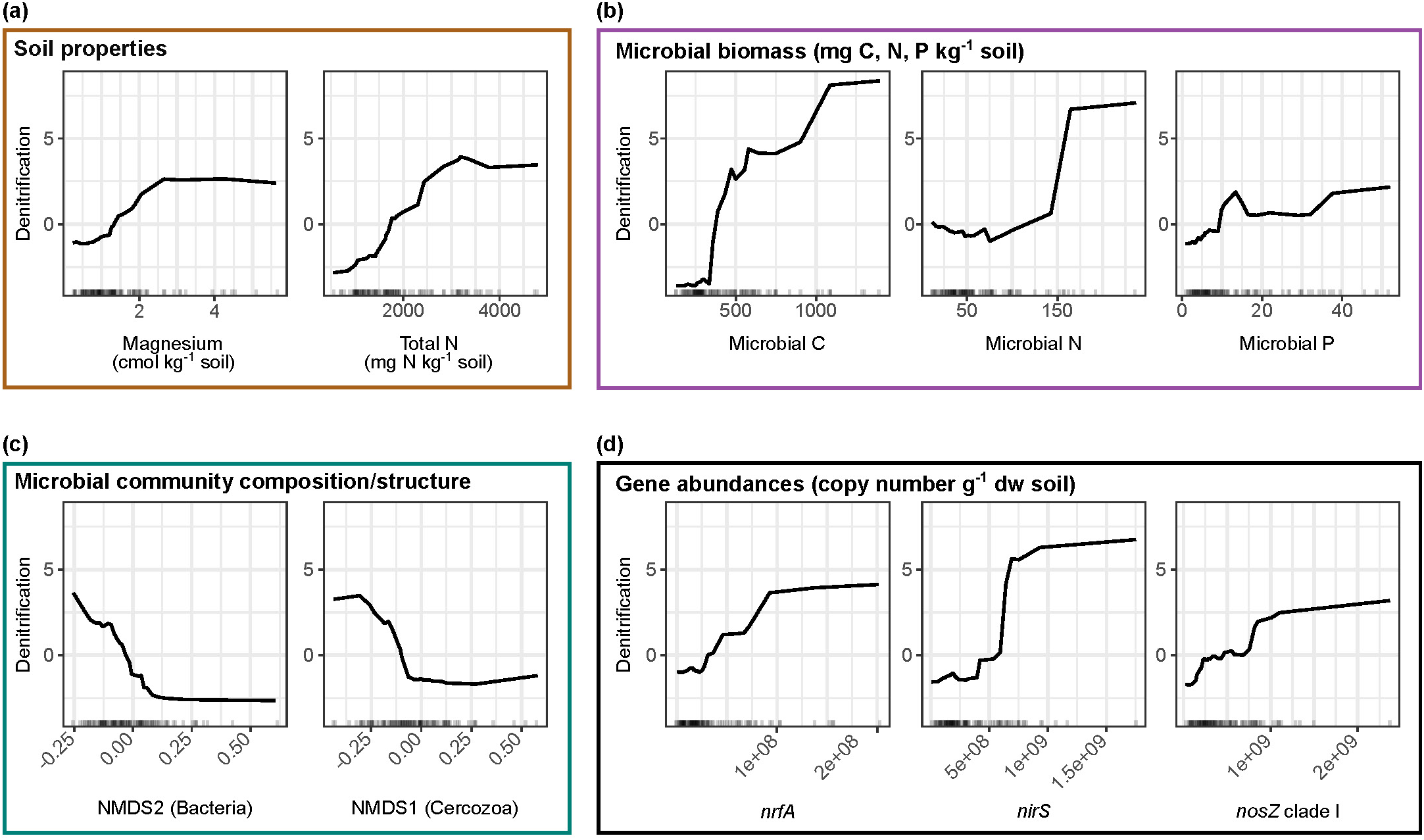
Relationships between potential denitrification activity (in ng N_2_O-N g^−1^ dw min^−1^) and its best predictors across European arable soils. Predictor variables selected by VSURF were used to generate accumulated local effects plots, which show the differences in prediction of denitrification compared to the mean prediction along the range of each predictor (x-axis), while accounting for potential correlations amongst predictor values. The effect is centred so that the mean effect is zero. The random forest model was built with 500 trees, 5 features considered at each split and a tree depth set to 6 (variance explained: 75 %, root mean square error: 69.9).

**Fig. 5.**
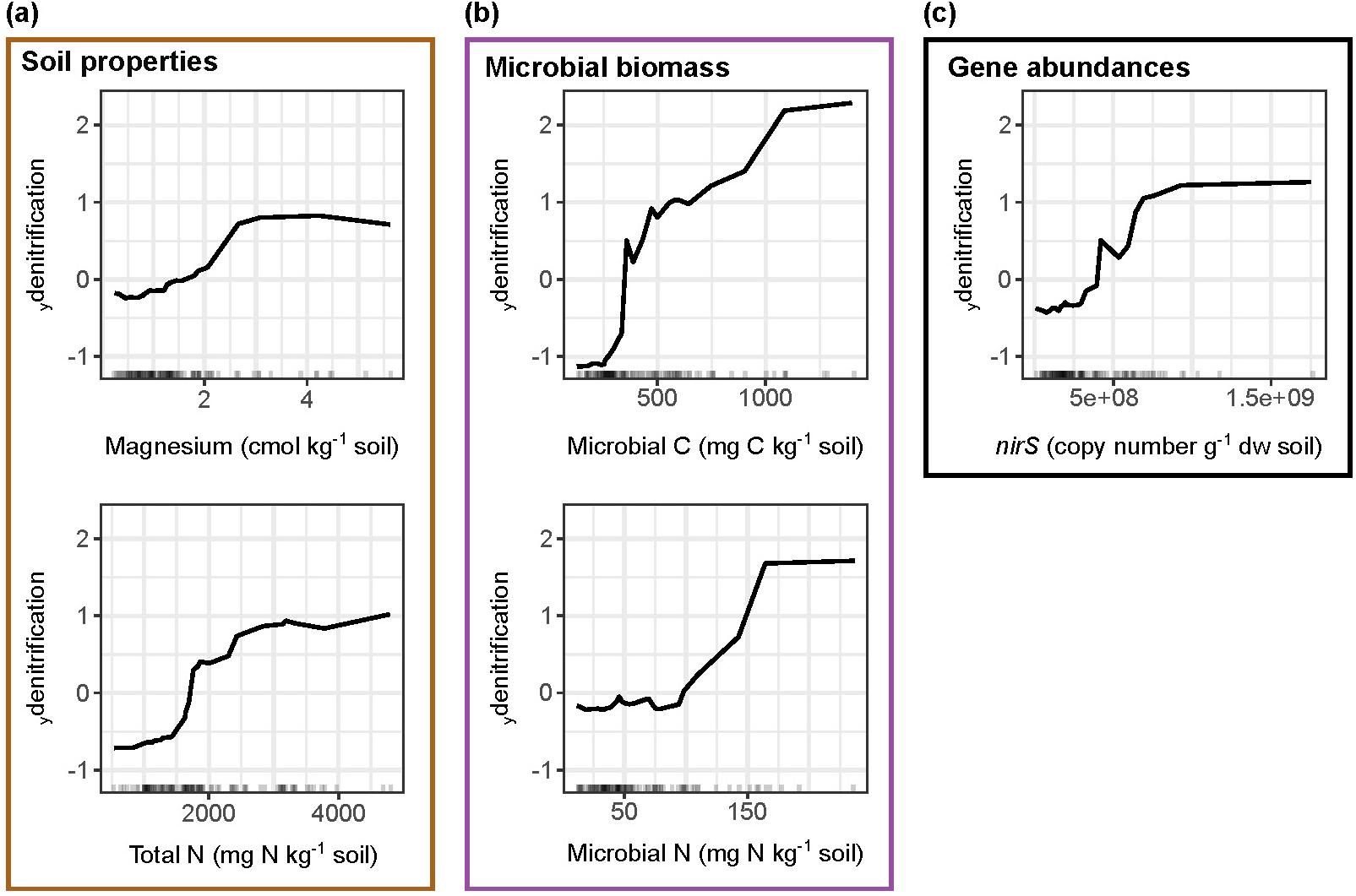
Relationships between yield-scaled potential denitrification activity (_y_denitrification = denitrification in ng N_2_O-N g^−1^ dw min^−1^/raw yield) and its best predictors across European arable soils. Predictor variables selected by VSURF were used to generate accumulated local effects plots, which show the differences in prediction of _y_denitrification (y-axis) compared to the mean prediction along the range of each predictor (x-axis), while accounting for potential correlations amongst predictor values. The effect is centred so that the mean effect is zero. The random forest model was built with 500 trees, 2 features considered at each split and a tree depth set to 9 (variance explained: 58 %, root mean square error: 4.3).

### Combining management and denitrification predictors

Finally, we assessed whether management practices could indirectly influence denitrification and _y_denitrification by affecting their best predictors. The redundancy analysis models, including the best predictors as response variables (**Figs. 4** and **5**) and the six management practices as explanatory variables, showed that management explained 37.5 % and 32.3 % of the variance in the predictors of denitrification and _y_denitrification, respectively (*p* = 0.001). Forward selection of the management variables in the full model identified three management practices as drivers, namely proportion of time with crop cover, application of organic fertilizers and tillage frequency, but their individual contribution was relatively small (3.0-5.5 %, **Fig. 6**). These practices correlated positively with all denitrification predictors, except the bacterial and cercozoan NMDS axes (**Fig. S1**). The inclusion of climate (mean annual temperature and mean annual precipitation over the past 10 years) and soil texture (clay and silt content) variables improved the models, which explained 56.9 % and 53.9 % of the variance in the predictors of denitrification and _y_denitrification, respectively (*p* = 0.001; **Fig. S2**). Although clay content and mean annual precipitation over the past 10 years were the two variables explaining most of the variation (9.3-12.3 %) in both models, the collective contribution of the different management, climate and soil texture variables was comparable (9-13 %).

**Fig. 6.**
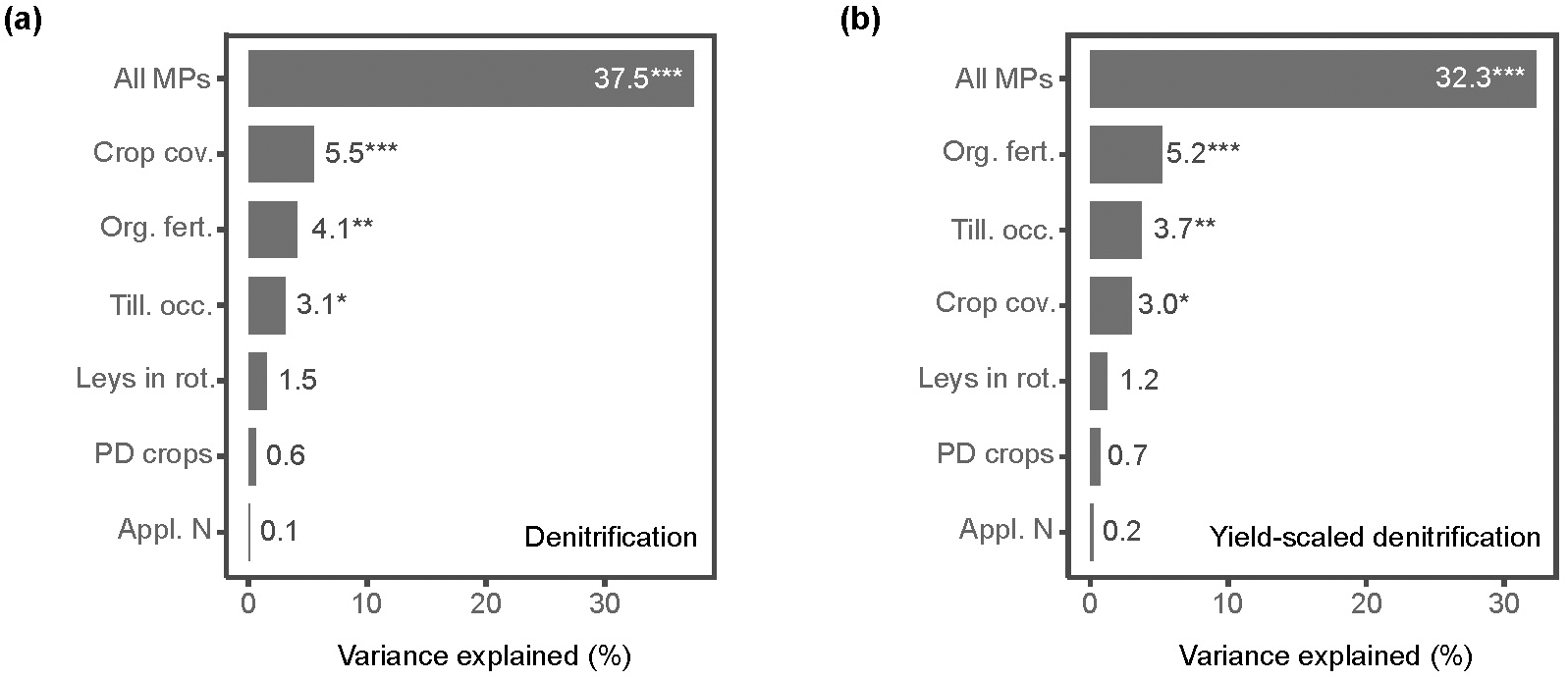
Variance explained by the full redundancy analysis model (all management practices, MP) and individual management practices after accounting for the remaining management practices in partial models for both (a) denitrification and (b) yield-scaled potential denitrification activity (_y_denitrification = denitrification/raw yield) predictors. Crop cov., proportion of time with crop cover (past 10 years); Org. fert., organic fertilizer application (past 4 years); Till. occ., tillage occurrences (sampling year); Leys in rot., presence of ley (past 10 years); PD crops, phylogenetic diversity of the crops (past 10 years); Appl. N, applied N fertilizer (sampling year).

## DISCUSSION

Increased time with crop cover has positive effects on soil biodiversity and functioning, including the accrual and accumulation of organic matter^14,21^ and the capacity to provide multiple ecosystems functions and services^12,14^. However, our study shows that crop cover also increases the risk for gaseous N losses through denitrification, which aligns with studies showing higher N_2_O emissions from fields with cover crops^22,23^. This is possibly explained by the increased carbon input through overall higher primary production combined with crop residue degradation leading to the formation of anoxic hotspots that favour denitrification as well as N_2_O emissions^24,25^. Higher crop diversity, measured here as the phylogenetic diversity of the crops in the rotations over 10 years prior to sampling, and fewer tillage occasions were associated with lower denitrification in relation to the harvested yield (_y_denitrification). Both practices have been shown to promote multiple ecosystem services, including soil fertility and carbon sequestration but differ in their influence on yield^14^. In contrast to reduced tillage, crop diversification can enhance cereal yields (as shown earlier with this dataset)^12^, particularly at low N fertilization rates^15,26^, and can also buffer against yield loss from global warming and changes in precipitation^27,28^. Moreover, diverse crop rotations can increase N-use efficiency in arable soils^29,30^, as supported here for _y_denitrification, thereby decreasing the amount of reactive N available for denitrification. In contrast to site and region specific reports indicating that cropping systems including perennials emit less N_2_O compared to systems with annual crops^31–33^, we found that the presence of leys in the rotations had no effect on denitrification, suggesting that the capacity of such rotations to decrease gaseous N losses from soils depends on a combination of the local environment and management. Overall, we conclude that agronomic management practices aiming to increase belowground diversity and support multiple ecosystems functions need to consider both possible trade-offs and co-benefits that may impact sustainability and productivity goals.

Denitrification was best predicted by the cumulative effect of N inputs, as reflected by total soil N, rather than the amount of N fertilizer applied in the sampling year. The growing surplus of reactive N in arable soils causes profound shifts in both the composition and functioning of soil microbial communities^34–36^, and this aligns with the acceleration of yearly increases in global N_2_O emission rates^37^. Surprisingly, there was no direct effect of addition of organic fertilizers during the past four years^38^. Nevertheless, organic fertilizer amendment affected the best predictors of both denitrification and _y_denitrification, indicating an indirect effect of organic amendments on denitrification. Apart from soil N content, several microbial variables, including microbial biomass, the composition of microbial communities and abundance of specific N-cycling guilds, predicted denitrification capacity across European croplands. The abundance of *nirS* and *nosZ* clade I, two genes indicating the presence of denitrifying microorganisms^39^, best explained effects of the functional guilds. We also found that denitrification was predicted by the abundance of nitrate ammonifiers (i.e., *nrfA*), likely because similar conditions are favourable for both denitrifiers and nitrate ammonifiers, even though nitrate ammonification leads to N retention rather than loss^40^. However, the genetic potential for nitrate ammonification was one to two orders of magnitude lower than that of denitrification (average *nrfA*:*nir* abundance ratio = 0.04±0.02). This suggests a limited mitigating effect of ammonification on N losses in arable soils across Europe and aligns with recent findings showing that denitrification capacity dominates over nitrate ammonification in terrestrial biomes^41^. Regarding the bacterial OTUs associated with an increase in the prediction of denitrification activity, the high diversity of physiological traits at the class level^42^ and the limited congruence between denitrification capacity and organismal phylogeny^43^ prevented further inferences on specific OTUs for denitrification capacity. The absence of variables related to fungal abundance and diversity in the set of the best denitrification predictors suggests that, relative to bacteria, denitrifying fungi^44^ contribute less to denitrification in arable soils. This could be because fungal denitrifiers make up a small proportion of the overall fungal community as well as of the total denitrifying community^45^. At any rate, the existence of a threshold for total N but not for gene abundances of N cycling guilds indicates that microbial factors are an important control of denitrification in arable soils. Future research on how resource availability shapes the interactions and niche partitioning within and between the different N-cycling guilds may provide additional insight into the mechanisms that link community composition and gaseous N losses in arable soils.

To conclude, our analysis of the relationships between denitrification and a set of 56 climatic, soil, microbial, and management variables associated with 106 agricultural soils across Europe revealed that denitrification activity was directly and indirectly influenced by management practices aiming to promote belowground diversity, particularly the proportion of time with crop cover. However, when denitrification was linked to crop yield by using yield-scaled denitrification, increased phylogenetic diversity of the crops in the rotations was associated with lower gaseous N losses. Taken together, our results thus support the widespread adoption of diverse crop rotations as a promising way to contribute to food security while minimizing negative impacts on the environment^16^.

## METHODS

### Sampling and associated data

The sampling procedure and part of the data used in this study have been described in detail in Garland et al.^12^. Briefly, soil samples were collected in fields under cereal cultivation across Europe (in Sweden, Germany, Switzerland, France and Spain) around flowering time between May and August 2017, depending on the country and site. In this study, we used a subset of the samples (n = 106) from Garland et al.^12^ for which information on management practices, crop yield, soil properties and microbial communities were available (**Fig. 1b-c**; **Table S1**). Management practices and crop yield were obtained by surveying the farmers and farm managers at each site via a questionnaire. Physical and chemical soil properties were measured using the Swiss standard protocols^46^. Potential rates for mineralization were determined following Gregorich and Carter^47^. Mean annual air temperature and precipitation (2007-2016) were obtained using the GPS coordinates of each sampling site and the historical monthly weather data from the WorldClim database (https://worldclim.org), using the R package ‘raster’ v. 3.6-26^ref.48^. A weighed measure of Faith’s phylogenetic diversity in the crop rotations was calculated using the weighted.faith function (https://rdrr.io/github/NGSwenson/lefse_0.5/) and a phylogeny generated with the package ‘V.PhyloMaker’ v. 0.1.0^ref.49^ in the R software v. 4.4.1^ref.50^.

OTU tables for archaea, bacteria, cercozoan, and fungi obtained from Garland et al.^12^ were rarefied by averaging the OTU counts over 1,000 computations using the rrarefy function in ‘vegan’ v. 2.6-4^ref.51^. The species abundance distribution patterns of the OTUs were then examined to partition the rarefied table between frequent and rare community members^52^. This was achieved by calculating an index of dispersion for each OTU, which corresponded to the ratio of the variance to the mean abundance multiplied by the occurrence^53^. The index was then used to model whether OTUs followed a stochastic (Poisson) distribution. Those falling below the 2.5% confidence limit of the *χ*^2^ distribution^54^ were discarded. Shannon’s diversity and Pielou’s evenness were determined using ‘vegan’ on the full rarefied tables with the diversity function. Non-metric multidimensional scaling (NMDS) ordinations were computed using the metaMDS function and Hellinger-transformed tables containing the frequent OTUs only. Focusing on the frequent community members minimizes the risk of sampling artefacts that can bias the distribution of the rare OTUs and thus increases the likelihood to detect relevant associations with the potential for denitrification.

### Potential denitrification activity

Potential denitrification activity was determined on soil samples kept at −20 °C using the acetylene inhibition technique modified from Pell et al.^55^. A slurry was prepared for each sample in 125 ml Duran bottles by adding 20 ml of water to 10 g fresh weight soil. The bottles were tightly capped, and the headspace was exchanged by flushing with N_2_ to obtain anoxic conditions. After 30 minutes of pre-incubation at 25 °C on a shaker (175 rpm), acetylene was added equivalent to 10% (v/v) of the headspace to inhibit the reduction of N_2_O to N_2_. Then, 1 ml of substrate was added to reach a final concentration of 3 mM KNO_3_, 1.5 mM succinate, 1 mM glucose, and 3 mM acetate^56^. Gas samples were taken from the headspace after 30, 75, 120, 150 and 180 minutes. Nitrous oxide concentrations were determined using a gas chromatograph (Clarus-500, Elite-Q PLOT phase capillary column, Perkin-Elmer, Hägersten, Sweden) equipped with a ^63^Ni electron-capture detector and the rate of N_2_O accumulation was determined in each bottle by non-linear regression.

We also adapted the widely used ‘yield-scaled N_2_O emissions’ metric (e.g.^ref.57^) to ‘yield-scaled N_2_ + N_2_O emissions’ (_y_denitrification), corresponding to the potential denitrification activity divided by the crop yield for the corresponding field, to determine whether any management practice can off-set denitrification activity with higher yields..

### Quantitative PCR analyses

The abundance of the 16S rRNA gene, fungal ITS and a set of nine genes involved in various steps in the N cycle (i.e. nitrogen fixation, nitrification, denitrification and ammonification; **Fig. 1b**) were measured by quantitative real-time PCR based on SYBR green detection. The qPCR reactions were carried out in duplicate runs on either the CFX Connect Real-Time System (Bio-Rad, Hercules, CA, USA) or ViiA7 (Life Technologies, Carlsbad, CA, USA) machine. Standard curves were obtained by serial dilutions of linearized plasmids with cloned fragments of the specific genes. The amplifications were validated by melting curve analyses and gel electrophoresis. Potential inhibition of PCR reactions was checked for all samples by amplifying a known amount of the pGEM-T plasmid (Promega, Madison, WI, USA) with the plasmid specific T7 and SP6 primers when added to the DNA extracts or non-template controls. No inhibition was detected with the amount of DNA used. Primer sequences and concentration, qPCR conditions and amplification efficiencies can be found in **Table S4**.

### Relationships between denitrification and management

Relationships between management practices and denitrification or _y_denitrification rates were assessed using generalized additive models. The models were fitted using the ‘mgcv’ package v. 1.9-1^ref.58^ with a gamma distribution for the response (denitrification or _y_denitrification) and the log link function. Fertilizer amount, phylogenic diversity of the crops and proportion of time with crop cover were modeled as thin plate splines (with *k* = 10) and tillage occurrences, application of organic fertilizer and presence of leys in the rotations as parametric terms. Country was included as a random factor. The gam.check function was used to ensure that enough basis functions (*k*) were specified for each smooth and that model residuals were normally distributed^58^. Concurvity between smooth terms, the non-linear equivalent of co-linearity, was assessed using the concurvity function implemented in the ‘mgcv’ package. Concurvity estimates were < 0.6 for all pair-wise comparisons. Model coefficients were estimated using restricted marginal likelihood^59,60^ and the goodness-of-fit of each model was calculated as the percentage of deviance explained. Effects were visualized on the response scale using the ‘visreg’ package v. 2.7.0^ref.61^.

### Identification of the predictors of denitrification activity

Random forest modelling was used to determine the relationships between denitrification and _y_denitrification and their best predictors among climatic, microbial, soil and management variables. The best predictors were identified by variable selection using the ‘VSURF’ package v. 1.2.0^ref.62^ with default parameters and denitrification or _y_denitrification as response variable. The algorithm was run 100 times and only the variables selected in more than 95% of the runs were retained. Then, a grid search was conducted to find the optimal combination of tuning parameters for the models containing the best predictors (n = 10 for denitrification and n = 5 for _y_denitrification): the number of variables to randomly sample as candidates at each split, the minimal number of samples within the terminal nodes and the fraction of samples to train the model on (with n = 500 trees; ‘randomForest’ package v. 4.7-1.1^ref.63^. The search was run 100 times and the combination of parameters corresponding to the best model fit (i.e. lowest out-of-bag root-mean-square error) was selected. The relationship between each variable and denitrification or _y_denitrification was visualized using accumulated local effects plots (grid.size = 25) implemented in the ‘iml’ package v. 0.11.3^ref.64^. These plots show how the prediction changes on average over the range of each individual explanatory variable, while accounting for potential correlations amongst explanatory variables^65^.

### Relationships between denitrification predictors and management, climate and soil texture

Redundancy analysis models were used to explore the relationships between management practices and denitrification and _y_denitrification predictors. We ran separate models for the two sets of predictors, using the ‘vegan’ package, and significance testing was done using permutation tests. Forward selection (ordiR2step function with default settings) and significance testing with permutation tests were applied to identify the management practices that were statistically significant for explaining the variation in the denitrification and _y_denitrification predictors. Partial models were implemented to test the amount of variation explained by each individual management practice (constrained variance) while accounting for the variation explained by the other management practices (conditioned variance). The same procedure was used to determine the relative importance of management, climate (mean annual temperature and mean annual precipitation over the past 10 years) and soil texture (silt and clay) on the variation in denitrification and _y_denitrification predictors.

## AUTHOR CONTRIBUTIONS

S.H., M.G.A.v.d.H, F.T.M., L.P. and M.C.R. initiated the study and planned the field work. A.S., S.B., F.D., A.E., P.G-P., G.G., C.H., D.S.P., and S.R. contributed to data collection. A.S. and M.E.S performed the analyses and A.S., M.E.S, G.V, R.B and S.H. interpreted the results. A.S., M.E.S and S.H. drafted the manuscript with input from G.V. and R.B. All authors commented on and approved the final manuscript.

## ACKNOWLEDGMENTS

The Digging Deeper project was funded through the 2015-2016 BiodivERsA call, with national funding from the Swiss National Science Foundation (grant 31BD30-172466), the Deutsche Forschungsgemeinschaft (grant 317895346), the Swedish Research Council Formas (grant 2016-0194 and 2018-02321), the Spanish Ministerio de Economía y Competitividad (grant PCIN-2016-028) and the Agence Nationale de la Recherche (grant ANR-16-EBI3-0004-01). We thank Claudia von Brömssen (Swedish University of Agricultural Sciences) for advice on the generalized additive models.

## DATA AVAILABILITY STATEMENT

The R code, OTU tables and metadata used in this study are available at Zenodo (10.5281/zenodo.14760398).

