## Supplementary information for "Diverse crop rotations off-set yield-scaled nitrogen losses via denitrification"

**Content:**

**Table S1.** Variables included in this study.
**Table S2.** Summary statistics of the GAM with denitrification as response variable.
**Table S3.** Summary statistics of the GAM with yield-scaled denitrification as response variable.
**Table S4.** qPCR conditions and amplification efficiencies.

**Figure S1.** Relationship between management practices and the best predictors for denitrification and yield-scaled potential denitrification activity.
**Figure S2.** Variance explained by the redundancy analysis models including management, climate and soil texture variables.

**Table S1.** List of variables included in this study and associated units

| Variable | Unit |
|---|---|
| **Management practices** | |
| Phylogenetic diversity of the crops (past 10 years) | / |
| Proportion of time with crop cover (past 10 years) | / |
| Presence of ley (past 10 years) | yes/no |
| Tillage occurrences (sampling year) | / |
| Applied N fertilizer (sampling year) | kg N $ha^{-1}$ |
| Organic fertilizer application (past 4 years) | yes/no |
| **Soil properties** | |
| Bulk density | kg $m^{-3}$ |
| Carbon:nitrogen ratio | / |
| Calcium content | cmol $kg^{-1}$ |
| Cation exchange capacity | cmol $kg^{-1}$ |
| Magnesium content | cmol $kg^{-1}$ |
| Moisture | g water $g^{-1}$ soil |
| Organic carbon | % |
| pH | / |
| Texture (silt & clay) | % |
| Total carbon | % |
| Total nitrogen | % |
| Total phosphorus | % |
| **Gene abundances** | |
| *nifH* | copies per g dry weight soil |
| Archaeal *amoA* | copies per g dry weight soil |
| Bacterial *amoA* | copies per g dry weight soil |
| *nxrB* (Nitrospira) | copies per g dry weight soil |
| *nirK* | copies per g dry weight soil |
| *nirS* | copies per g dry weight soil |
| *nosZ* clade I | copies per g dry weight soil |
| nosZ clade II | copies per g dry weight soil |
| *nrfA* | copies per g dry weight soil |
| *nosZ/nir* | / |
| *nrfA/nir* | / |
| 16S rRNA gene | copies per g dry weight soil |
| Fungal ITS | copies per g dry weight soil |
| **Microbial biomass** | |
| Microbial C | mg C $kg^{-1}$ soil |
| Microbial N | mg N $kg^{-1}$ soil |
| Microbial P | mg P $kg^{-1}$ soil |
| **$\alpha$- and $\beta$-diversity (Archaea, Bacteria, Cercozoa and Fungi*)*** | |
| Shannon's diversity | / |
| Pielou's evenness | / |
| NMDS1 | / |
| NMDS2 | / |
| **Activity rates** | |
| Potential mineralization | mg N $kg^{-1}$ soil $day^{-1}$ |
| Potential denitrification activity | ng $N_20$-N $g^{-1}$ soil dw $min^{-1}$ |
| Yield-scaled potential denitrification activity | ng $N_20$-N $g^{-1}$ soil dw $min^{-1}$ $t^{-1}$ $ha^{-1}$ |
| **Climate** | |
| Mean annual temperature (2007-2016) | °C |
| Mean annual precipitation (2007-2016) | mm |

**Table S2**. Summary of the generalized additive model (GAM) used to assess the effects of management practices on the potential denitrification activity of the soils (n = 106). Explanatory variables included phylogenetic diversity of the crops (past 10 years; pd), proportion of time with crop cover (past 10 years; pCC), presence of ley in the rotations (past 10 years), amount of fertilizer applied (sampling year; N_fert), application of organic fertilizer (past 4 years) and tillage occurrences (sampling year). Country was included as a random factor.

| **Smooth terms** | **Ref edf** | **edf** | **F statistic** | ***p*-value** |
|---|---|---|---|---|
| f(pd) | 1.00 | 1.00 | 2.57 | 0.11 |
| f(pCC) | 3.58 | 2.87 | 2.75 | 0.03 |
| f(N_fert) | 2.03 | 1.62 | 1.00 | 0.38 |
| Country | 4.00 | 3.35 | 7.87 | 1.92e-06 |
| **Parametric coefficients** | **Estimate** | **Std. error** | | ***p*-value** |
| Tillage occ. (2-3) | 0.12 | 0.21 | | 0.54 |
| Tillage occ. (4-6) | 0.26 | 0.29 | | 0.36 |
| Appl. organic fert. | 0.24 | 0.24 | | 0.32 |
| Presence of leys | -0.01 | 0.19 | | 0.94 |
| Intercept coefficient: 2.12 ± 0.32 | | | | |
| Deviance explained: 47.3 % | | | | |

**Table S3**. Summary of the generalized additive model (GAM) used to assess the effects of management practices on the yield-scaled potential denitrification activity of the soils (n = 106). Explanatory variables included phylogenetic diversity of the crops (past 10 years; pd), proportion of time with crop cover (past 10 years; pCC), presence of ley in the rotations (past 10 years), amount of fertilizer applied (sampling year; N_fert), application of organic fertilizer (past 4 years) and tillage occurrences (sampling year). Country was included as a random factor.

| **Smooth terms** | **Ref edf** | **edf** | **F statistic** | ***p*-value** |
|---|---|---|---|---|
| f(pd) | 1.00 | 1.00 | 3.93 | 0.05 |
| f(pCC) | 2.18 | 1.73 | 0.58 | 0.54 |
| f(N_fert) | 1.00 | 1.00 | 0.37 | 0.55 |
| Country | 4.00 | 3.29 | 5.44 | 9.37e-05 |
| **Parametric coefficients** | **Estimate** | **Std. error** | | ***p*-value** |
| Tillage occ. (2-3) | 0.46 | 0.24 | | 0.05 |
| Tillage occ. (4-6) | 0.77 | 0.34 | | 0.02 |
| Appl. organic fert. | 0.23 | 0.28 | | 0.41 |
| Presence of leys | 0.01 | 0.22 | | 0.95 |
| Intercept coefficient: 0.24 ± 0.36 | | | | |
| Deviance explained: 37.6 % | | | | |

**Table S4.** Primers, thermal cycling conditions and amplification efficiency for quantification of nitrogen cycling genes.

| **Process** | **Gene**<br>Primer names | **Sequences** (5'-3') | **Conc. (µM)** | **Thermal cycling** | **Efficiency** |
|---|---|---|---|---|---|
| | **Fungal ITS**[1] | | | (95°C, 5 min) x 1 | |
| | ITS3F | GCATCGATGAAGAACGCAGC | | (95°C, 15s, 55°C 30s, 72°C 30s, 78°C 5s) x 35 | 88 % |
| | ITS4R | TCCTCCGCTTATTGATATGC | | *(95°C, 15 s; (60 to 95° C, 5 s, increment 0.5°)), x 1* | |
| | **16S rRNA**[2] | | | (95°C, 5 min) x 1 | |
| | 341F | CCTACGGGAGGCAGCAG | 0.5 | (95°C, 15s, 60°C 30s, 72°C 30s, 78°C 5s) x 35 | 105 % |
| | 534R | ATTACCGCGGCTGCTGGCA | 0.5 | *(95°C, 15 s; (60 to 95° C, 5 s, increment 0.5°)), x 1* | |
| **$N_2$ fixation** | ***nifH***[3] | | | (95°C 5min) x 1 | |
| | Po 1F | TGCGAYCCSAARGCBGACTC | 0.5 | (95°C 15s, 55°C 30s, 72°C 30s, 80°C 10s) x 35 | 98-105 % |
| | Po 1R | ATBGCCATCATYTCRCCGGA | 0.5 | *(95°C, 15 s; (60 to 95° C, 5 s, increment 0.5°)), x 1* | |
| **Ammonia ox.** | ***amoA***[4] | | | (95°C, 5 min) x 1 | |
| (archaeal) | crenamoA23F | ATGGTCTGGCTWAGACG | 0.5 | (95°C, 15 s; 55 °C, 30 s; 72°C, 40 s; 77°C, 5 s) x 40 | 85% |
| | crenamoA616R | GCCATCCATCTGTATGTCCA | 0.5 | *(95°C, 15 s;(60 to 95° C, 5 s, increment 0.5°)), x 1* | |
| **Ammonia ox.** | ***amoA***[5] | | | (95°C, 5 min) x 1 | |
| (bacteria) | AmoA1F | GGGGTTTCTACTGGTGGT | 0.5 | (95°C, 15s, 55°C 30s, 72°C 40s, 80°C 5s) x 35 | 83 % |
| | AmoA2R | CCCCTCKGSAAAGCCTTCTTC | 0.5 | *(95°C, 15 s; (60 to 95° C, 5 s, increment 0.5°)), x 1* | |
| **Nitrite ox.** | ***nxrB***[6] | | | (95°C 5 min) x 1 | |
| (*Nitrospira*) | nxrB169f | TACATGTGGTGGAACA | 0.5 | (95°C 15s, 56°C 30s, 72°C 45s, 78°C 8s) x 35 | 89-98 % |
| | nxrB638r | CGGTTCTGGTCRATCA | 0.5 | *(95°C, 15 s; (60 to 95° C, 5 s, increment 0.5°)), x 1* | |
| **Denitrification** | ***nirK***[7] | | | (95°C, 5 min) x 1 | |
| Nitrite | 876F | ATYGGCGGVAYGGCGA | 0.5 | (95°C, 15 s; (63°C – 58°C, -1°/cycle), 30 s; 72°C, 30 s) x 6 | 100 % |
| reduction | 1040R | GCCTCGATCAGRTTRTGGTT | 0.5 | (95°C, 15 s; 58°C, 30 s; 72°C, 30 s; 80°C, 5 s) x 35 | |
| | | | | *(95°C, 15 s; (60 to 95° C, 5 s, increment 0.5°)), x 1* | |
| **Denitrification** | ***nirS***[8] | | | (95°C, 5 min) x 1 | |
| Nitrite | S4QF | GTSAACGYSAAGGARACSGG | 0.5 | (95°C, 15 s; (65°C – 60°C, -1°/cycle), 30 s; 72°C, 35 s) x 6 | 88 % |
| reduction | S6QR | GASTTCGGRTGSGTCTTSAYGAA | 0.5 | (95°C, 15 s; 58°C, 30 s; 72°C, 35 s; 80°C, 5 s) x 35 | |
| | | | | *(95°C, 15 s; (60 to 95° C, 5 s, increment 0.5°)), x 1* | |
| **Denitrification** | ***nosZI***[9] | | | (95°C, 7 min) x 1 | |
| Nitrous oxide | 1840F | CGCRACGGCAASAAGGTSMSSGT | 0.8 | (95°C, 15 s; (65°C – 60°C, -1°/cycle), 30 s; 72°C, 30 s) x6 | 97 % |
| reduction | 2090R | CAKRTGCAKSGCRTGGCAGAA | 0.8 | (95°C, 15 s; 60°C, 30 s; 72°C, 30 s; 80°C, 5 s) x 35 | |
| | | | | *(95°C, 15 s; (60 to 95° C, 10 s, increment 0.5°)), x 1* | |

| | | | | | |
|---|---|---|---|---|---|
| **Denitrification** | ***nosZII***[10] | | | (95°C, 7 min) x 1 | |
| Nitrous oxide | nosZII-F | CTIGGICCIYTKCAYAC | 0.8 | (95°C, 15 s; 54°C, 30 s; 72°C, 30 s; 77°C, 5 s) x 40 | 79 % |
| reduction | nosZII-R | GCIGARCARAAITCBGTRC | 0.8 | (95°C, 15 s; (60 to 95° C, 10 s, increment 0.5°)), x 1 | |
| **Ammonification** | ***nrfA***[11] | | | (95°C, 5 min) x 1 | |
| Nitrite | nrfAF2aw | CARTGYCAYGTBGARTA | 0.5 | (95°C, 15 s; (57°C – 52°C, -1°/cycle), 30 s; 72°C, 30 s) x6 | 91-105 % |
| reduction | nrfAR1 | TWNGGCATRTGRCARTC | 0.5 | (95°C, 15 s; 52°C, 30 s; 72°C, 30 s; 80°C, 10 s) x 39 | |
| | | | | *(95°C, 15 s; (60 to 95° C, 5 s, increment 0.5°)), x 1* | |

[1]: White *et al*. 1990; [2]: Muyzer *et al*. 1993; [3]: Poly *et al*. 2001; [4]: Tourna *et al*. 2008; [5]: Rotthauwe *et al*. 1997; [6]: Pester *et al*. 2004; [7]: Henry *et al*. 2004; [8]: Kandeler *et al*. 2006; [9]: Henry *et al*. 2004; [10]: Jones *et al.* 2013; [11]: Cannon *et al*. 2019.

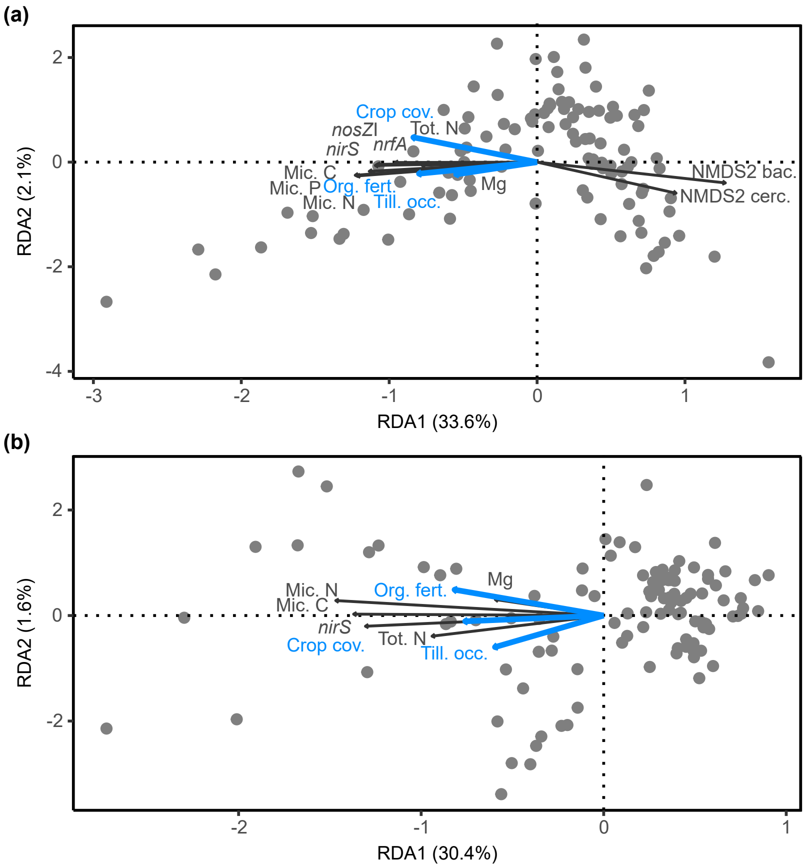


**Figure S1.** Relationship between management practices and the best predictors for denitrification and yield-scaled potential denitrification activity ($_y$denitrification = denitrification/raw yield). Redundancy analysis plots show the direction of relationships between the significant management practices and the best predictors of (a) denitrification and (b) $_y$denitrification. Sites are depicted by grey dots and the arrows indicate denitrification predictors (black arrows) and management practices (blue arrows). Mic. C, N and P, Microbial carbon, nitrogen and phosphorus; Mg, soil magnesium; Tot. N, total soil nitrogen; Crop cov., proportion of time with crop cover (past 10 years); Org. fert., organic fertilizer application (past 4 years); Till. occ., tillage occurrences (sampling year).

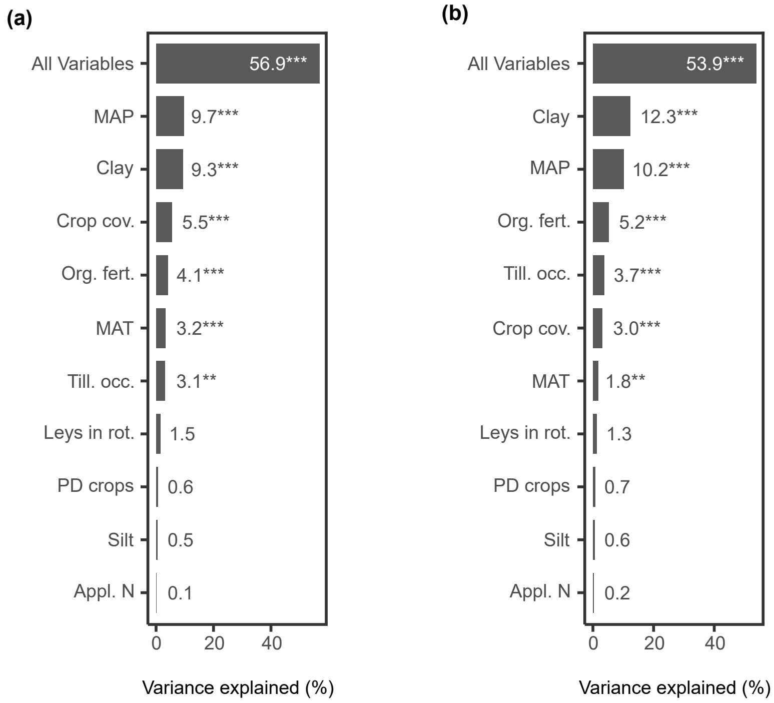


**Figure S2.** Variance explained by the full redundancy analysis model (all variables, i.e. management practices, climate data and soil texture) and individual variables after accounting for the remaining variables in partial models for both (a) denitrification and (b) yield-scaled potential denitrification activity (ydenitrification = denitrification/raw yield) predictors. Crop cov., proportion of time with crop cover (past 10 years); Org. fert., organic fertilizer application (past 4 years); Till. occ., tillage occurrences (sampling year); Leys in rot., presence of ley (past 10 years); PD crops, phylogenetic diversity of the crops (past 10 years); Appl. N, applied N fertilizer (sampling year); MAP, mean annual precipitation (2007-2016); MAT, mean annual temperature (2007-2016); Clay, soil clay content (%); Silt, soil silt content (%).
